# Encoding and Retrieval in Parallel: ERP Correlates of Continuous Recognition Memory for Natural Scenes

**DOI:** 10.64898/2026.07.07.736108

**Authors:** Elena Cesnaite, Niko A. Busch

**Affiliations:** Institute of Psychology, University of Münster, Germany; Otto-Creutzfeldt-Center for Cognitive and Behavioral Neuroscience, University of Münster, Germany

## Abstract

Human long-term memory for visual scenes is remarkably robust, yet the neural mechanisms supporting memory encoding and retrieval remain poorly understood when both processes must operate at the same time. For instance, this might happen when we encounter a familiar place while simultaneously forming new memories of this encounter. We investigated electrophysiological correlates of visual recognition memory using a continuous recognition task (CRT), in which participants judged a continuous stream of scene photographs as previously seen or new, such that encoding and retrieval occurred in parallel on every trial. To make recognition particularly demanding, stimuli were drawn from only four scene categories. Thirty-one participants performed the task while EEG was recorded, and we analyzed canonical ERP markers of retrieval (mid-frontal FN400, 300-550 ms; late parietal effect, LPE, 550-800 ms) and encoding (subsequent memory effect, SME) as a function of stimulus repetition and lag between consecutive presentations. FN400 showed robust old/new effects for both repetitions, whereas LPE differences emerged only at the second repetition. While FN400 amplitude was insensitive to lag, LPE amplitude decreased systematically with increasing lag, mirroring the behavioral pattern of declining accuracy and slower responses. A significant SME emerged selectively for images subsequently recognized on both repetitions, indicating that the SME in continuous recognition is specific for the most robustly encoded items and reflects the strength of encoding. Together, these findings show that canonical ERP markers of recognition memory are preserved even when encoding and retrieval operate concurrently, but their expression depends on how often and how recently an item has previously been encoded – parameters that can be flexibly manipulated within the CRT. This demonstrates that the CRT is sensitive to fine-grained temporal dynamics of memory formation and retrieval that could be missed under standard single-repetition designs.

## 1 Introduction

Human memory for visual scenes is strikingly robust. Pioneering studies have shown that people can remember thousands of images with high accuracy, even after long delays (Brady et al., 2008; Shepard, 1967). Notably, this visual memory is not only long-lasting but also resilient to interference, even when individuals are exposed to many exemplars from the same scene category (Konkle et al., 2010).

To better understand the cognitive and neural mechanisms that support such high-capacity visual memory, recognition memory has been extensively investigated using event-related potentials (ERPs). This research has enabled the identification of several well-characterized ERP components associated with memory processes (Voss & Paller, 2017; Wilding & Ranganath, 2011). During encoding, the subsequent memory effect (SME) reflects differential neural activity for items that are later remembered compared to those that are forgotten. SME has been found in recall and recognition tasks, and presumably reflects several cognitive processes that lead to more efficient encoding, such as detailed perceptual analysis, rote rehearsal, semantic elaboration, or mental imagery (Voss & Paller, 2017).

During retrieval, early and late ERP old/new effects are typically observed. The early frontal old/new effect, often referred to as the FN400 (approximately 300–500 ms post-stimulus), is characterized by more negative amplitudes for new relative to old items at frontal electrodes. Within dual-process theories of recognition memory, this effect has been linked to familiarity-based recognition, reflecting a graded sense that an item has been encountered before without retrieval of contextual details. A later parietal old/new effect (often referred to as LPE), typically emerging between 500 and 800 ms, has been more strongly associated with recollection, that is, the retrieval of episodic details about the prior encounter. This late parietal positivity is commonly interpreted as indexing the conscious reinstatement of contextual information accompanying successful memory retrieval (Curran, 2000; Curran & Cleary, 2003; Liesefeld et al., 2016).

Most ERP studies of recognition memory have used an experimental paradigm with two clearly separated stages: an initial encoding phase, during which a set of novel stimuli is presented for later memory, followed by a retrieval phase, during which participants indicate if they recognize an item as old (previously seen) or new. However, such a strict temporal separation between encoding and retrieval phases does not align well with everyday memory, where both processes often occur simultaneously. For example, during a daily commute, encountering a familiar place may trigger a recognition response (“I’ve been here before”) while simultaneously updating or strengthening the already-formed memory trace of that place. Similarly, a novel stimulus such as a newly built construction site may be categorized as new while it is being encoded for future reference. With repeated exposure, this previously novel location becomes increasingly familiar. These examples highlight that encoding and retrieval are not always distinct phases but can occur concurrently.

To capture this continuous mode of memory processing, researchers have employed the continuous recognition task (CRT; Fox & Osth, 2023; Hockley, 1982; Shepard & Teghtsoonian, 1961). In this paradigm, stimuli are presented in a continuous sequence, and participants indicate for each item whether it is new or has been seen before. Crucially, this task structure requires participants to engage in both encoding and retrieval on every trial, rather than being in a dedicated “encoding mode” or “retrieval mode” as in conventional two-phase paradigms. The CRT affords several important methodological and theoretical advantages. First, because participants must both retrieve old items and encode new ones throughout the task, the CRT more closely mirrors the dynamics of natural memory processing. Second, the strength of memory traces can be systematically manipulated via the lag between repetitions, i.e. by varying the number of intervening items between the first and second presentation. This manipulation affects both temporal delay and the degree of interference from intervening stimuli. Third, because there is no dedicated retention interval between encoding and test, there is minimal opportunity for rehearsal or strategic reorganization of the material (Shepard & Teghtsoonian, 1961). Importantly, varying lag also enables a distinction between recognition based on short-term or working memory (e.g., at short lags within a few seconds) and long-term memory (at longer lags), providing a unique window into the temporal dynamics of memory consolidation and forgetting (Friedman, 1990). Finally, the CRT offers a pragmatic advantage in terms of experimental design flexibility. Unlike standard paradigms that depend on the completion of both an encoding and a subsequent retrieval phase, the CRT allows for continuous data collection. Old and new items are interleaved throughout the session, enabling the analysis of both encoding and retrieval processes at any point in the stream. This flexibility makes the CRT especially attractive for studies with variable session lengths, such as online experiments investigating visual long-term memory (sometimes referred to as a “memory game”; Bylinskii et al., 2015; Goetschalckx & Wagemans, 2019; Kramer et al., 2023).

Compared to the large body of research using paradigms with separate encoding and retrieval phases, relatively few ERP studies have employed the CRT to investigate the neural correlates of visual recognition memory. In a seminal study, Friedman, 1990 presented line drawings of everyday objects within a CRT paradigm and reported ERP old/new effects that resembled the canonical early frontal and late parietal components typically observed in standard recognition tasks. Interestingly, the magnitude of these ERP effects was unaffected by the lag between an item’s initial and repeated presentation. This insensitivity to lag could suggest that the observed old/new effects reflect the outcome of a binary recognition decision – classifying an item as either old or new – rather than a graded signal of memory strength. Alternatively, the stimulus set, comprising only a single exemplar per object category, may have been highly memorable, and the maximum lag of 32 intervening items may not have been sufficient to induce substantial forgetting (Nickerson, 1965). This interpretation is supported by the observation that behavioral performance remained stable even at longer lags.

Moreover, Friedman, 1990 found no evidence for subsequent memory effects (SMEs). The absence of SME in a CRT task may imply that SMEs only emerge when cognitive resources can be fully devoted to encoding, i.e. in a dedicated encoding phase without the competing demands of simultaneous retrieval. Another possibility is that the lack of SMEs resulted from the high memorability of the stimuli, which may have required little elaboration or strategic encoding to be successfully remembered (Voss & Paller, 2017). Taken together, these findings leave open the question whether the neurocognitive mechanisms that support visual recognition and encoding during continuous, concurrent processing resemble those observed under conditions of strictly separated encoding and retrieval.

The present study aimed to investigate the neural correlates of visual recognition memory under continuous, ecologically valid conditions. Specifically, we examined whether conventional ERP memory effects, such as the SME during encoding and old/new effects during retrieval, can be observed in a CRT using realistic natural scene photographs, where discrimination between old and new items is intentionally made challenging by presenting numerous exemplars from the same scene categories. In addition, we assessed how SME encoding effects evolve across repeated presentations of the same item, thereby probing the dynamics of encoding processes under ongoing memory demands. Finally, we investigated how the ERP old/new effect varies as a function of five lag conditions between stimulus repetitions to better understand how memory strength and retrieval processes unfold over time in the context of continuous recognition.

## 2 Methods

### 2.1 Participants

Data were collected from 33 participants (22 female; 4 left-handed, mean age 27 years, SD 5.5 years) with normal or corrected-to-normal vision^1^. All participants provided written informed consent, and received course credit or monetary compensation. The study was approved by the ethics committee of the faculty of psychology and sports sciences, University of Münster (approval no. 2016-22-N).

### 2.2 Stimuli and apparatus

The experiment was written in MATLAB (The Mathworks, Natick, MA, USA) using the Psychophysics Toolbox (Brainard, 1997). Stimuli were presented on a calibrated LCD monitor (VIEWPixx/EEG) with 1920 x 1080 pixels resolution and 120 Hz refresh rate, placed at a distance of 86 cm from the participants’ eyes. Head position was stabilized using a chin rest.

The stimuli were drawn from a pool of 1200 images from 4 different scene categories (forests, highways, beaches, cities) gathered using Google Image Search. All images were 10 x 7.5 *^◦^* in size and presented in grayscale on black background. A gray fixation cross was displayed at the center of the screen during the inter-stimulus intervals, turning red or green for 200 ms after the response to provide feedback.

**Figure 1:**
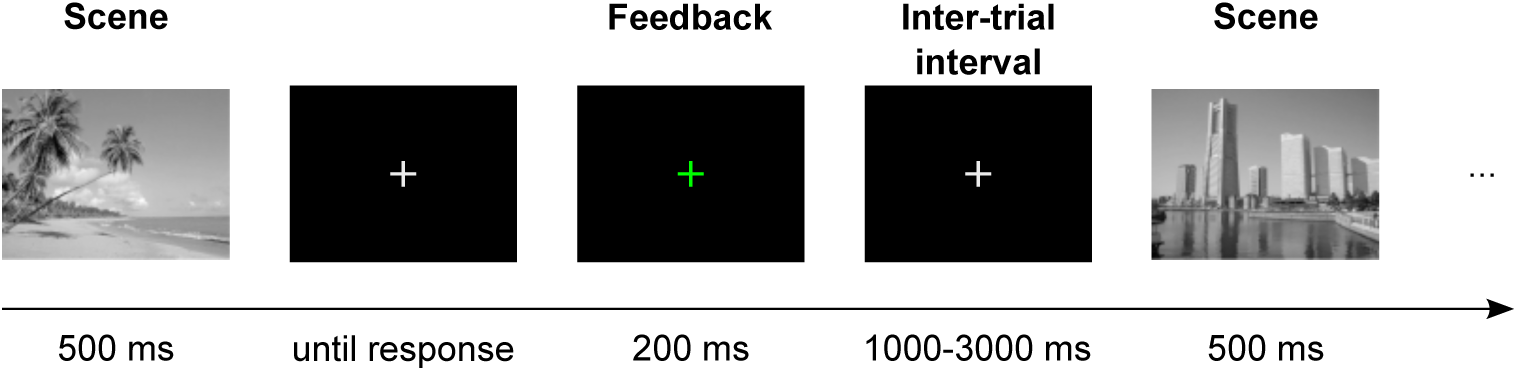
Timeline of an example trial in the EEG continuous recognition memory task. Participants held down two keys to initiate a trial. Scene images (beaches, forests, highways, or cities) were presented for 500 ms. Participants indicated whether they had seen the image before by releasing one of the keys as quickly as possible. Feedback was provided via a change in fixation cross color (green = correct, red = incorrect).

### 2.3 Procedure

In this CRT (Friedman, 1990; Shepard & Teghtsoonian, 1961), participants were presented with a stream of images (Figure 1), some of which were repeated, and they decided for each image whether it had been presented before (old) or whether it was the first time that image was presented (new). Participants were instructed to respond as fast as possible by lifting their finger from one of two response keys (left/right CTRL key), indicating whether the image was old or new. The assignment of key (left/right) and response category (old/new) was counterbalanced across participants. Images were presented for 500 ms each, followed by an empty screen until the participant had responded. Feedback was provided by turning the fixation cross green or red for correct and incorrect responses, respectively, for a duration of 200 ms. Once both keys were held down again, the next image followed after a variable inter-trial interval, which was drawn from a truncated exponential distribution with a minimum of 1000 ms and a maximum of 3000 ms.

For each participant, a sample of 600 images was selected from the image pool; 300 were presented only once and 300 images were presented three times (first presentation as a new image, second and third presentations as old), resulting in a total of 1200 trials per participant, half of which featured a new image. Repeated presentations occurred after a lag of 10, 16, 24, 38, or 60 intervening trials.

### 2.4 EEG Recording

Data were recorded at two laboratory locations in sound-attenuated rooms. One laboratory location (17 participants) was equipped with a Faraday cage, whereas the other location (16 participants) had no electrical shielding. In both labs, EEG was recorded with a BioSemi Active-Two amplifier system from 64 Ag/AgCl electrodes arranged according to the international 10-10 system and two additional mastoid electrodes. The horizontal and vertical electro-oculograms were recorded from additional electrodes at the lateral canthi of both eyes and below the eyes, respectively. Two additional electrodes located adjacent to electrode POz served as reference and ground. Signals were sampled at 1024 Hz with a 200 Hz low-pass filter.

### 2.5 EEG preprocessing

EEG data were re-referenced to the ‘Cz’ channel, low-pass filtered at 40 Hz (Blackman-windowed FIR filter, filter order 282, 10 Hz transition bandwidth), and high-pass filtered at 0.1 Hz (Hamming-windowed FIR filter, filter order 846, 2 Hz transition bandwidth). EEG data were then downsampled to 256 Hz.

Data segments containing gross artifacts (i.e., muscle artifacts) were rejected by manual selection. Broken or unresponsive channels were identified by visual inspection of power-spectral density and removed from the data. Ocular and other noise artifacts were corrected using extended infomax independent component analysis (ICA). Given that ICA is sensitive to high-amplitude, low-frequency amplitude fluctuations (e.g., sweat artifacts), ICA was applied to optimized training data, created by additionally high-pass filtering the data at 1 Hz. We used the IClabel toolbox (Pion-Tonachini et al., 2019) to help identify independent components associated with eye movement, muscle, heart, and channel noise artifacts. Minimal classification accuracy was set at 0.9. The resulting unmixing weights were then transferred to the original (i.e., less severely high-pass filtered) dataset. The strong high-pass filter was not applied to the original data to avoid the loss of potentially cognition-related signals (see Dimigen, 2020; Tanner et al., 2016; Widmann et al., 2015).

We then interpolated missing channels by using spherical spline interpolation and re-referenced data to a common average reference. Finally, the continuous EEG data were segmented into epochs from -200 to 800 ms relative to stimulus onset and baseline-corrected using the -100 to 0 ms pre-stimulus interval.

Based on previous literature, analyses of old/new and lag effects focused on data from correct trials in two predefined time windows and corresponding regions of interest. ERP amplitudes were averaged across frontal electrodes (Fz, F1, F2, AFz) in the 300-550 ms time window, and across parietal electrodes (Pz, P1, P2, CPz) in the 550-800 ms time window. These early frontal and late parietal selections are commonly used to quantify the FN400 and LPE old/new effects, respectively (Curran & Cleary, 2003; Voss & Paller, 2017). The analysis of the SME focused on frontal electrodes (Fz, F1, F2, AFz) in the 300-550 ms time window and parietal electrodes (PO7, PO3, PO8, PO4) in the 550-800 ms time window. Single-trial amplitude values below Q1 - 1.5 × the interquartile range (IQR) or above Q3 + 1.5 × IQR were considered outliers and removed from subsequent analyses.

### 2.6 Participant exclusion

Two participants were excluded from further analyses due to chance-level behavioral performance. One participant’s accuracy was only 53.7%, 46.6%, and 51.9% for the first, second, and third stimulus presentations, respectively. The second participant showed low accuracy on the first presentation (39.6%) but higher accuracy on the subsequent presentations (66% and 70%), suggesting a response bias toward indicating that stimuli had been seen before, which may have artificially inflated performance on later presentations.

### 2.7 Statistical analyses of ERP and behavioral data

Behavioral performance (mean accuracy and response times) was compared between image presentations (first/new, second, third) and lags (10, 16, 24, 38, 60) using separate linear models in R (lm function). Note that we could not use lag and presentation within a single model because lag is not defined for the first presentation. The models were specified as:

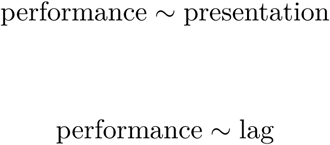

FN400 and LPE mean amplitudes in the early frontal and late parietal regions of interest were compared between image presentations and lags using linear mixed-effects models implemented in R (lme4 package). Presentation and lag conditions were included as fixed effects, and a random intercept for participant was specified to account for inter-individual variability. The models were specified as:

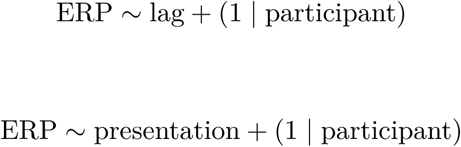

The SME at frontal and parietal areas was analyzed using a linear mixed-effects model. ERP amplitudes at the first presentation were predicted by subsequent memory performance, defined as the number of correct recognitions across the second and third presentations (0, 1, or 2). The model included subsequent memory as a fixed effect and a random intercept for participant and was specified as:

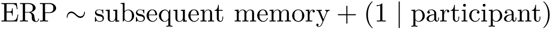

## 3 Results

### 3.1 Behavioral results

**Figure 2:**
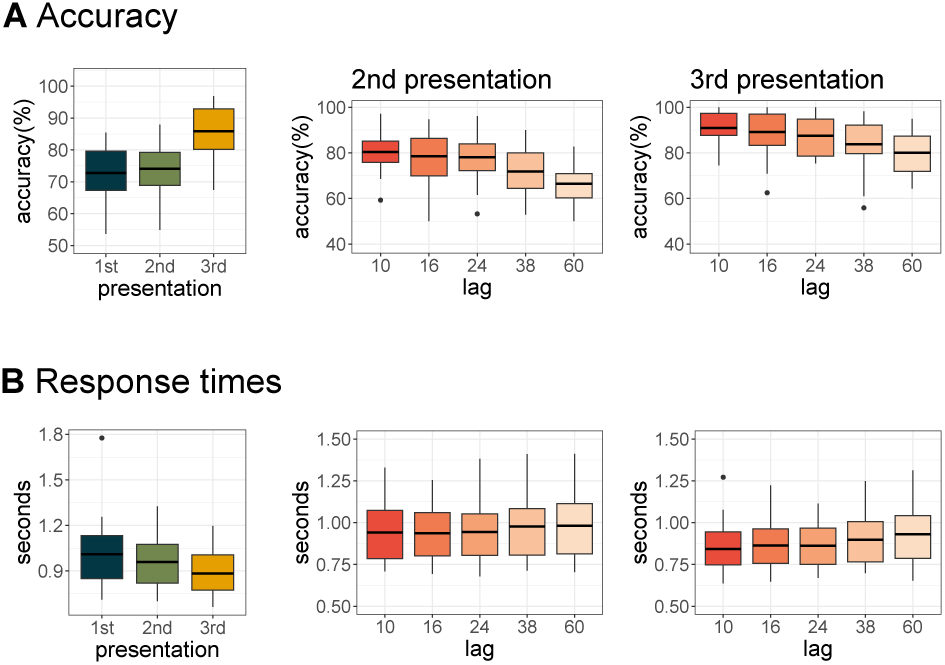
**A**: Recognition accuracy. Left: accuracy for scenes at first (new), second, and third (old) presentations, averaged across all lags. Middle: accuracy for second presentations (old) as a function of lag since the previous presentation. Right: accuracy for third presentations (old) as a function of lag. **B**: Response times (same conventions as in A).

We used linear mixed-effects models to test whether recognition memory accuracy and response times differed as a function of presentation and lag.

Accuracy and response times did not differ between the first and second presentation (accuracy: *b* = 1.3, *p* = 0.53; *RT* : *b* = −0.05*, p* = 0.25). In contrast, the third presentation yielded higher accuracy and faster responses than the first presentation (accuracy: *b* = 13.1, *p <* 0.0001; RT: *b* = −0.13, *p* = 0.0059; Figure 2A).

Accuracy decreased linearly with increasing lag for both the second (*b* = −3.46, *p <* 0.0001) and third presentations (*b* = −2.73, *p <* 0.0001). Response times were not significantly related to lag for the second presentation (*b* = 0.012, *p* = 0.221), but they increased as a function of lag in the third presentation (*b* = 0.02, *p* = 0.013; Figure 2B).

### 3.2 ERP results

#### Old/new effects

At frontal channels, the ERP waveforms showed a large negative deflection peaking within the conventional FN400 time window (300–550 ms; Figure 3 A). Within this time window, frontal ERPs were more negative for new images (first presentation) than for old images (as compared to the second presentation: *b* = 0.55, *p <* 0.0001; and as compared to the third presentation: *b* = 0.44, *p <* 0.0001), consistent with a FN400 old/new effect.

At parietal channels, the ERP waveforms showed a large positive deflection extending across the 550–800 ms time window (Figure 3 B). There was no significant difference between the first and second presentations of an image (*b* = 0.11, *p* = 0.27) on LPE amplitude at parietal channels. However, LPE amplitude was significantly more positive for old images at the third presentation as compared to new images (*b* = 0.87, *p <* 0.0001).

#### Lag effects

At frontal channels, there was no statistically significant relationship between the amplitude of FN400 component and lag conditions (*b* = 0.02, *p* = 0.72; Figure 3C).

At parietal channels, LPE showed a significant negative amplitude shift with increasing lag (*b* = −0.13, *p* = 0.006; Figure 3D). That is, the positive parietal deflection was systematically attenuated with greater lag between consecutive presentations.

#### Subsequent memory effects

At frontal channels, FN400 showed no significant amplitude difference between new images that were subsequently never recognized and those subsequently recognized at only one of the two subsequent presentations (*b* = 0.43, *p* = 0.13; Figure 4A). However, new images that were subsequently recognized at both subsequent presentations elicited a more positive-going frontal response compared to never-recognized images (*b* = 0.72, *p* = 0.012; Figure 4A and B), reflecting an attenuation of the early frontal negativity. Specifically, amplitudes were on average 0.72 µV more positive for subsequently twice-recognized images than for never-recognized images.

At parietal channels, LPE likewise showed no significant amplitude difference between new images subsequently never recognized and those subsequently recognized only once (*b* = −0.28, *p* = 0.27). However, new images subsequently recognized at both following presentations elicited a less positive-going parietal response compared to never-recognized images (*b* = −0.64, *p* = 0.013; Figure 4C and D), reflecting an attenuation of the late parietal positivity. Specifically, amplitudes were on average 0.64 µV smaller for subsequently twice-recognized images than for never-recognized images.

**Figure 3:**
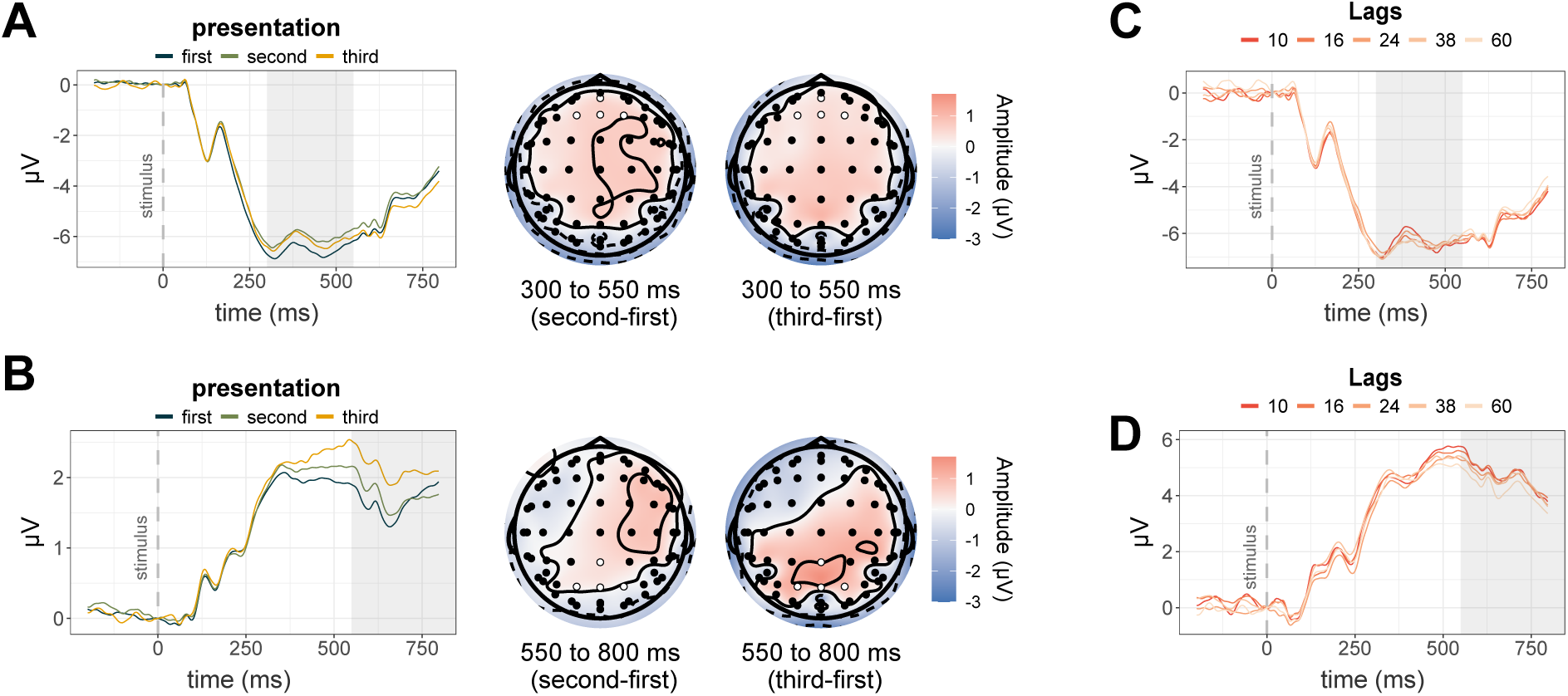
Grand-average ERP waveforms and scalp topographies of repetition and lag effects for scene photographs. **(A, B)** ERPs for the first (new; dark teal), second (light green), and third (yellow) image presentations at frontal (A) and parietal (B) electrode clusters, with corresponding scalp topographies showing the difference in mean amplitude between repeated (second or third) and first presentations during the FN400 (300–550 ms; A) and LPE (550–800 ms; B) time windows. **(C, D)** ERPs for five lag conditions (10, 16, 24, 38, and 60 intervening items; dark to light orange) at frontal (C) and parietal (D) electrode clusters. Shaded areas mark the analysis epochs of interest (300–550 ms for the FN400; 550–800 ms for the LPE). Electrodes included in the frontal and parietal clusters are marked in white.

**Figure 4:**
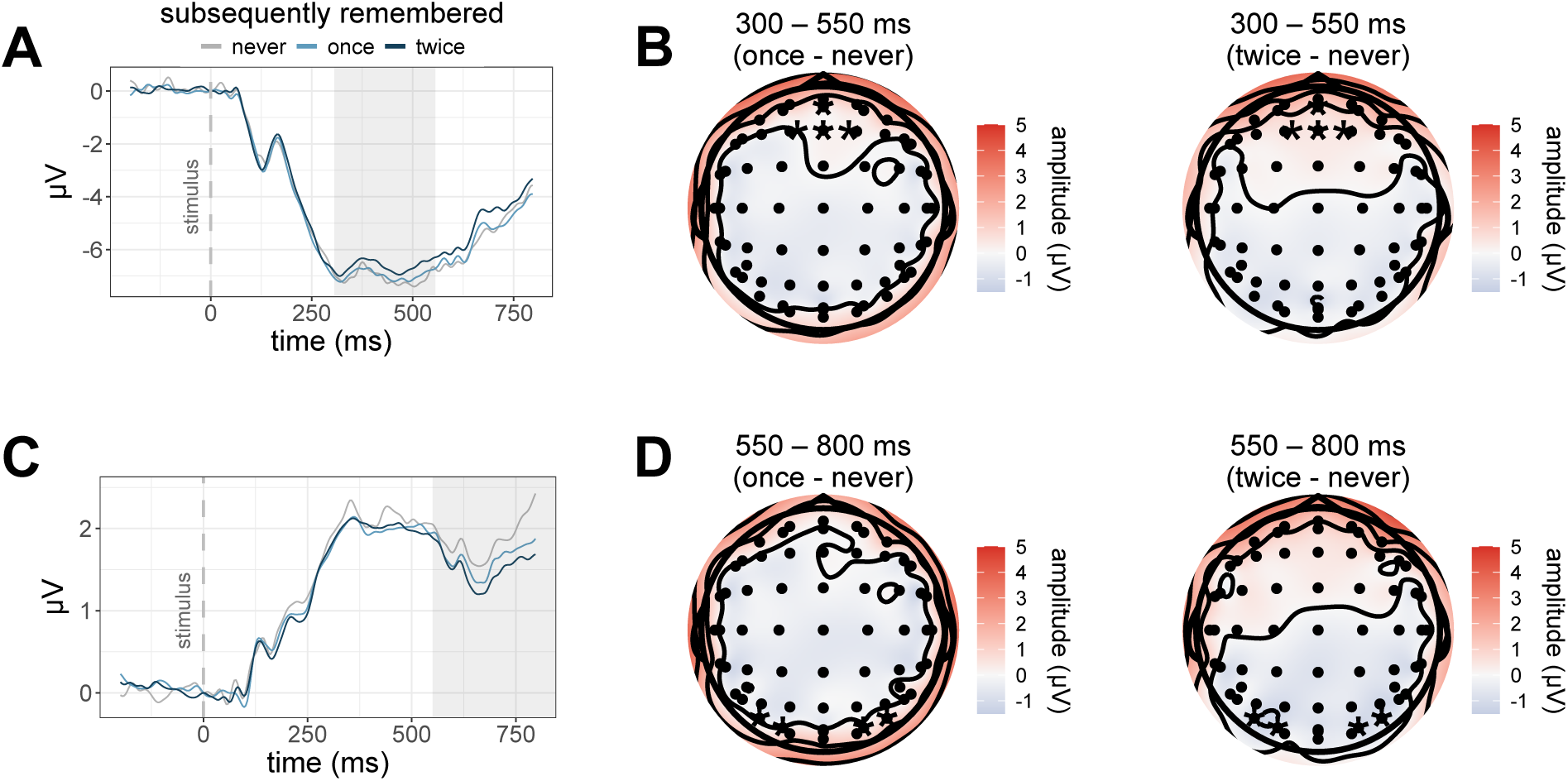
Grand-average ERP waveforms and scalp topographies of the subsequent memory effect (SME) for scene photographs. ERPs are shown separately for images that were subsequently never remembered (grey), remembered only once (light blue), or remembered twice (dark blue) at **(A)** frontal and **(C)** parietal electrode clusters. Shaded time windows indicate the analysis epochs of interest (300 to 550 ms and 550 to 800 ms). Scalp topographies in panels B and D display the difference in mean amplitude between subsequently remembered (once or twice) and never-remembered images, separately for the two time windows. Warm colors indicate greater positivity for subsequently remembered relative to never-remembered images. Electrodes included in the frontal and parietal clusters are marked with asterisks.

## 4 Discussion

How do the neurocognitive mechanisms of memory encoding and retrieval operate when both processes must occur in parallel, rather than in dedicated sequential phases? The continuous recognition task (CRT) offers a unique window into this question, as participants must simultaneously encode novel stimuli and retrieve previously seen ones on every trial (Shepard & Teghtsoonian, 1961). Only few ERP studies have employed this task (see Ellmore et al., 2022), and the one study most directly comparable to ours – Friedman (1990), using line drawings of objects – reported two unexpected null results: no subsequent memory effect (SME) and no modulation of old/new ERP effects by lag (i.e., the number of images in between consecutive presentations). These findings raised the possibility that the neurocognitive processes supporting encoding and retrieval are fundamentally altered when both demands compete concurrently for cognitive resources. The present study tested this hypothesis using naturalistic scene photographs, a wider range of lags, and multiple image repetitions, enabling a more sensitive assessment of encoding strength.

### Old/new effects

The ERP old/new effect is a widely studied phenomenon in recognition memory research characterized by more positive-going waveforms for old compared to new items. This effect is not a single unitary component but is comprised of several spatio-temporally distinct subcomponents that correlate with different mnemonic processes (Friedman & Johnson Jr., 2000; Voss & Paller, 2017).

We observed robust old/new ERP effects across stimulus repetitions: both FN400 (300–550 ms, frontal) and LPE (550–800 ms, parietal) showed a positive shift in amplitude for old compared to new items. While the frontal effect did not distinguish how often an image had been repeated, the parietal old/new effect only emerged for images presented for the third time, but not yet for images shown for the second time. A similarly graded shift across multiple repetitions has been reported by Johnson et al. (1998). These findings confirm that canonical ERP markers of recognition memory are preserved under continuous recognition conditions, replicating and extending previous studies (Berman et al., 1991; Friedman, 1990) to naturalistic scene photographs.

Within the dual-process framework of recognition memory (Curran, 2000; Curran & Cleary, 2003), the FN400 is broadly associated with familiarity-based recognition – a relatively automatic sense of prior encounter – while the LPE is thought to reflect recollection, that is, the conscious reinstatement of contextual details from the encoding episode (Wilding & Ranganath, 2011). While we did not assess familiarity and recollection directly, the progressive increase across presentations indicates that both familiarity and recollection are successfully recruited in the CRT, even as participants must simultaneously encode new items on each trial. This stands in contrast to the theoretical possibility that the dual demands of the CRT might selectively impair the more resource-intensive recollection process while sparing automatic familiarity.

The differential sensitivity of FN400 and LPE to presentation number also aligns with the behavioral data. While accuracy and response times improved primarily at the third presentation, FN400 amplitude was significantly larger at both the second and third presentations relative to the first. By contrast, the LPE effect emerged only at the third, closely tracking behavioral improvement. This is consistent with the FN400 reflecting an early, coarse familiarity signal that does not strengthen with further repetition, while the LPE indexes a more graded recollection-based process that builds with additional encoding opportunities. Taken together with the selective SME for items recognized on both repetitions, these findings suggest that especially the third presentation marks a shift toward a more robust, recollection-supported memory representation.

The interpretation of the FN400 as a neural correlate of familiarity-based recognition, as proposed by dual-process accounts (Curran, 2000; Curran & Cleary, 2003), has been contested in the literature. An influential alternative view holds that the FN400 reflects conceptual implicit memory or fluency – a facilitated processing of previously encountered conceptual information – rather than episodic familiarity in the strict sense (Rugg & Curran, 2007). Accordingly, the FN400 old/new effect would not necessarily indicate that a participant consciously recognizes an item as previously seen, but rather that its conceptual representation is processed more fluently upon repetition. This distinction is particularly relevant here, since stimuli belonged to only four scene categories, raising the possibility that our FN400 effect partly reflects category-level fluency rather than item-level episodic familiarity. However, because new items belonged to the same four categories as old items throughout the experiment, category-level fluency should have been equally present for both and thus cannot account for the old/new difference. Nevertheless, we acknowledge that the precise functional interpretation of the FN400 remains debated.

It is worth noting that recognition in the present study was considerably more demanding than in most prior ERP work, because participants had to discriminate between numerous exemplars from the same scene category rather than items from distinct conceptual categories, probably resulting in substantial categorical interference (Konkle et al., 2010). The robustness of old/new ERP effects under these conditions underscores the sensitivity of both FN400 and LPE as markers of visual recognition memory even when conceptual identity cannot serve as a reliable retrieval cue.

### Lag effects

A further novel finding concerns the differential sensitivity of the FN400 and LPE to the lag between consecutive image presentations, which is an indicator of memorial decay (Berman et al., 1991), possibly due to item noise and context drift (Fox & Osth, 2023). While FN400 amplitude did not vary as a function of lag, LPE amplitude decreased systematically with increasing lag, mirroring the behavioral pattern of declining accuracy and slower responses at longer lags. This dissociation suggests that familiarity-based recognition is relatively robust to the number of intervening items and the interference they introduce, while recollection-based retrieval is more sensitive to memory strength and degrades more steeply with increasing lag.

Notably, Friedman (1990) observed no modulation of either the FN400-like or LPE-like old/new effects across lags. As argued in the Introduction, this may have reflected a ceiling effect arising from the high memorability of distinctive object exemplars and a relatively short maximum lag of 32 intervening items. By employing a broader lag range (10 to 60 intervening trials) and stimuli more susceptible to forgetting due to within-category similarity, our design might have been better positioned to detect lag-dependent modulations.

### Subsequent memory effects

The most theoretically significant finding of the present study is the observation of a significant subsequent memory effect (SME) in the CRT with images, in direct contrast to Friedman (1990). Critically, the SME emerged only for images remembered on *both* subsequent presentations, not for images remembered only once. At the first presentation, images later recognized twice elicited greater FN400 positivity and greater LPE negativity compared to images never subsequently recognized, whereas no significant ERP difference was observed between once-remembered and never-remembered images.

Items encoded only weakly – sufficient for one later recognition but not two – may not have engaged elaborative encoding processes strongly enough to produce detectable ERP differences relative to forgotten items. Only images encoded robustly enough to support consistent recognition across two subsequent encounters showed a reliable SME. This interpretation is in line with the view that the SME indexes the depth or elaborateness of initial encoding (Voss & Paller, 2017), and suggests that under the competing demands of the CRT, only the most thoroughly encoded items leave a detectable neural trace at encoding.

The absence of an SME in Friedman (1990) is thus likely attributable, at least in part, to a methodological constraint: with only a single repetition per image, subsequent memory could only be classified as remembered or forgotten without distinguishing between weakly and strongly encoded items. Our two-repetition design revealed that the SME is not absent in continuous recognition, but is selective for robustly encoded stimuli.

Notably, while the positive FN400 SME at frontal electrodes is consistent with the direction most commonly reported in the literature (Voss & Paller, 2017), the SME at parietal electrodes showed the opposite polarity, with more negative amplitudes for subsequently remembered items. Mecklinger and Kamp (2023) proposed that the direction of the SME depends on the nature of the encoding process engaged: positive SMEs are associated with semantic elaboration, whereas negative SMEs reflect reliance on perceptual feature processing. Interestingly, studies using targeted manipulations of the encoding task with word stimuli have generally found a consistently positive or negative SME across electrode sites, rather than a combination of both polarities (Bridger & Wilding, 2010; Otten & Rugg, 2001). The simultaneous presence of both polarities in our data may reflect the particular demands of our task, which required discriminating between numerous exemplars drawn from only four scene categories (beaches, forests, highways, cities). Because within-category exemplars share the same semantic identity, successful subsequent recognition could not rely on semantic attributes alone and instead required additional encoding of perceptual details (“road scene with traffic jam and a yellow truck”). This dual encoding demand may explain the coexistence of both SME polarities, whereas studies using more easily discriminable stimuli, where a single encoding strategy may suffice, typically report a single (usually positive) polarity (Duarte et al., 2004).

### Limitations

Several limitations of the present study warrant consideration. First, our stimulus set was restricted to four scene categories. While this restriction was theoretically motivated – to impose within-category discrimination demands – it limits the generalizability of the findings to other scene categories or other stimuli such as faces or objects. Second, the CRT by design does not permit full separation of encoding- and retrieval-related ERP activity, since both processes co-occur on every trial. While this is an inherent constraint of the paradigm, it is simultaneously one of its ecological virtues, as it more closely mirrors the conditions under which memory operates in everyday life.

## Conclusion

The present study demonstrates that the neurocognitive mechanisms supporting visual recognition memory are not fundamentally disrupted when encoding and retrieval must occur in parallel in a continuous recognition task. Canonical ERP markers of recognition memory – the FN400 and the LPE – were robustly observed for naturalistic scene photographs, with amplitudes scaling with the number of repetitions. Critically, a lag-dependent dissociation between the two components revealed that recollection, but not familiarity, is sensitive to memory strength in the context of continuous recognition. Most importantly, we identified a SME that was selective for images subsequently remembered on both repetitions, reconciling the previously reported absence of the SME in CRT paradigms with the rich SME literature from two-phase designs. Together, these findings reveal that the CRT is not merely a pragmatic alternative to conventional paradigms, but a sensitive tool for probing the concurrent dynamics of memory encoding and retrieval under ecologically valid conditions.

## Data and Code Availability

Code for data processing, analysis, and figures can be found at https://github.com/ecesnaite/ CORENATS. The data and stimuli can be found at https://doi.org/10.17879/w7rr8-yym45 and https://doi.org/10.17879/pnqjm-w5s77, respectively.

## Author Contributions

**Conceptualization:** Elena Cesnaite and Niko A. Busch. **Data curation:** Elena Cesnaite and Niko A. Busch. **Formal analysis:** Elena Cesnaite and Niko A. Busch. **Funding acquisition:** Niko A. Busch. **Investigation:** Elena Cesnaite and Niko A. Busch. **Methodology:** Elena Cesnaite and Niko A. Busch. **Project administration:** Elena Cesnaite and Niko A. Busch. **Resources:** Niko A. Busch. **Software:** Elena Cesnaite and Niko A. Busch. **Supervision:** Niko A. Busch. **Visualization:** Elena Cesnaite. **Writing:** Elena Cesnaite and Niko A. Busch.

## Footnotes

1 This dataset was additionally used in the EEGManyPipelines project to investigate variability in EEG analysis practices (Cesnaite et al., 2025, 2026; Trübutschek et al., 2024). These meta-scientific questions are independent of the memory-related research questions addressed in the present study.

